# Breeding and identification of promising Mauritius x Kew pineapple hybrids with high heterosis for fruit and plant traits

**DOI:** 10.1101/2023.10.09.561547

**Authors:** Lalit Dhurve, K. Ajith Kumar, Jyothi Bhaskar, A. Sobhana, Rose Mary Francies, Deepu Mathew

**Affiliations:** Department of Fruit Science, Kerala Agricultural University, Thrissur – 680 656, India; Centre for Plant Biotechnology and Molecular Biology, College of Agriculture, Kerala Agricultural University, Thrissur – 680 656, India; Regional Agricultural Research Station, Kerala Agricultural University, Ambalavayal, Wayanad – 673 593, India; Fruit Crops Research Station, Kerala Agricultural University, Thrissur – 680 656, India; Agricultural Research Station, Kerala Agricultural University, Mannuthy, Thrissur – 680 651, India

**Keywords:** *Ananas comosus*, Hybridization, Heterobeltiosis, Selection index, Standard heterosis, Variety evaluation

## Abstract

Leading cultivars of pineapple Mauritius and Kew were hybridized and 25 hybrids were evaluated under open field conditions, using randomized block design with two replications. Performance of the female parent cum check variety Mauritius, male parent Kew and check variety in Kerala state, India, Amritha, were also evaluated and compared. Based on the performance, heterobeltiosis, average heterosis and standard heterosis over two check varieties, in each hybrid, for 10 plant growth traits and 24 fruit traits, were calculated. For fruit weight, hybrid H35 had the highest heterobeltiosis and standard heterosis over Amritha whereas H62 had highest standard heterosis over Mauritius and average heterosis. For pulp weight, hybrid H17 had the highest values for all heterosis parameters. For TSS, hybrid H62 had the highest heterobeltiosis and other parameters were highest in H43. For days to attain physiological maturity, crown weight, peel weight and acidity, H27, H30, H77 and H43, respectively were lowest in all heterosis parameters. Based on the selection criterion [∑average heterosis (fruit weight, TSS, pulp weight) – ∑average heterosis (crown weight, peel weight, eye profile, eye relative surface, time taken for physiological maturity, acidity)] developed using the average heterosis values for desirable and undesirable fruit traits, six hybrids H66, H17, H59, H43, H70 and H35 were identified for further evaluation. The identified hybrids also satisfied the requirements in fruit weight (≥1.0 kg), pulp weight (≥750.58 g), TSS (≥14.49 ^○^Brix), days to attain physiological maturity (≤185.70 days), crown weight (≤305.50 g), peel weight (≤159.27 g) and acidity (≤1.05).

## Introduction

Pineapple (*Ananas comosus* L., Bromeliaceae) originated in southeastern Brazil, is cultivated widely in all the tropical and subtropical regions of the world (Cabot, 1992; Firoozabady and Gutterson, 2003). Pineapple cultivation in south India dates back to 1550 (Laufer, 1929) and in the state of Kerala itself, this crop covers 9,200 ha with cultivar ‘Mauritius’ occupying 95 per cent area (Ecostatkerala, 2020).

With lesser number of cultivars dominating, a wide genetic variability is missing in this crop. The fresh fruit market demands superior fruit characters including high TSS, medium size and longer post-harvest life whereas processing industry prefers Kew variety having bigger, cylindrical, juicy fruits with flat eyes. Even with this sort of specific demands, the crop improvement works in this crop has been significantly lower compared to other commercial fruits. So far, Amritha is the only pineapple hybrid released in India and even after having high TSS, this hybrid suffers from its average fruit size. Thus, it has become essential to hybridize and generate new lines combining desirable traits to fresh fruit as well as processing markets.

Mauritius with medium fruit size, high TSS, low acidity and high pulp recovery is the most preferred table variety whereas Kew with higher fruit size and juice content is good for processing (Dhurve et al., 2021). Crossing of these cultivars was done with the objective to identify the lines with larger cylindrical fruits with high TSS, low acidity and high pulp recovery. Even though no precise selection criterion has been defined in this crop, an accession yielding 1.5-2.5 times higher fruit weight along with other commercial parameters may be considered promising (Rasmusson, 1987; Brat et al., 2004). Thus, this study had the objective to evaluate the field level heterosis parameters for the plant growth and yield traits in the Mauritius x Kew hybrids and to select the best hybrid using a simple selection criterion formulated based on the fruit quality traits.

## Material and Methods

The hybridization of Mauritius and Kew, with Mauritius as female parent, was done at Fruit Crops Research Station (FCRS), Kerala Agricultural University, Thrissur, India. Cultivars of both the types, widely grown in the Kerala state, India, were used in the hybridization. Agronomic information on the parent cultivars and the check varieties used in the study are presented in Table 1.

**Table 1.** The agronomic information on the parents and the check varieties used in the study.

| Sl. No. | Trait | Mauritius | Kew | Amritha |
| --- | --- | --- | --- | --- |
| 1 | Plant height (cm) | 100.53 | 109.02 | 91.36 |
| 2 | Leaves per plant | 42.10 | 41.54 | 40.64 |
| 3 | Length of D leaf (cm) | 61.94 | 57.57 | 52.44 |
| 4 | Breadth of D leaf (cm) | 2.92 | 2.64 | 2.64 |
| 5 | D leaf area (cm<sup>2</sup>) | 130.97 | 109.97 | 100.13 |
| 6 | Leaf Area Index | 4.09 | 3.38 | 3.01 |
| 7 | Days to attain ideal leaf stage for flowering | 196.30 | 175.10 | 180.57 |
| 8 | Days for initiation of flowering (visual) | 48.90 | 39.74 | 49.40 |
| 9 | Days for 50 % flowering | 55.44 | 45.50 | 55.64 |
| 10 | Flowering phase (days) | 26.14 | 23.04 | 24.97 |
| 11 | Length of the fruit (cm) | 15.05 | 12.75 | 11.65 |
| 12 | Girth of the fruit (cm) | 31.90 | 34.10 | 30.00 |
| 13 | Breadth of the fruit (cm) | 8.31 | 9.47 | 8.34 |
| 14 | Crown weight (kg) | 0.10 | 0.21 | 0.10 |
| 15 | Taper ratio of the fruit | 0.83 | 0.84 | 0.85 |
| 16 | Peel weight (kg) | 0.15 | 0.14 | 0.13 |
| 17 | Pulp weight (kg) | 0.73 | 0.75 | 0.40 |
| 18 | Pulp percentage | 78.43 | 82.02 | 72.96 |
| 19 | Days for fruit maturity (days) | 140.77 | 138.00 | 140.87 |
| 20 | Crop duration (days) | 385.97 | 352.84 | 370.84 |
| 21 | Shelf life (days) | 7.94 | 7.50 | 7.25 |
| 22 | Juice (%) | 89.79 | 86.54 | 87.87 |
| 23 | TSS (° Brix) | 13.52 | 12.47 | 16.82 |
| 24 | Acidity (%) | 0.89 | 0.88 | 0.86 |
| 25 | Total sugars (%) | 11.99 | 11.69 | 10.86 |
| 26 | Reducing sugars (%) | 2.16 | 4.10 | 3.28 |
| 27 | Non-reducing sugars (%) | 9.83 | 7.59 | 7.58 |
| 28 | Sugar/ acid ratio | 15.19 | 14.17 | 19.56 |
| 29 | Fibre (%) | 33.06 | 35.10 | 34.71 |
| 30 | Total carotenoids (%) | 228.33 | 255.42 | 267.55 |
| 31 | Ascorbic acid (mg/ 100 g) | 64.79 | 44.27 | 50.09 |
| 32 | Fruit weight with crown (kg) | 1.02 | 1.13 | 0.65 |
| 33 | Profile of eyes | 4.80 | 4.00 | 5.00 |
| 34 | Relative surface of eyes | 4.47 | 5.00 | 5.00 |

Suckers of Mauritius and Kew were planted in open field under two-line system and managed following the standard procedures (KAU, 2016). At 39-42 leaf stage ethrel application was done and hybridization was started when the first flower in the inflorescence opened. Due to the self-incompatibility in pineapple, emasculation of the female flowers was not required before pollination. Flowers were isolated from the Kew inflorescence by making three deep triangular incisions into the base of the flower, at anthesis time (6.00-10.00 am). Pollen collected from the excised flowers stayed fresh longer with lower dehydration. For hand-pollination, anther was picked up with forceps, pollens were accumulated in the forceps tip by piercing the anther walls and gently brushed onto the stigma of opened Mauritius flowers. Contamination with foreign pollen was prevented by covering the pollinated inflorescence using a paper bag. Only three to seven flowers of the 100-200 flowers in an inflorescence open every day, and hence two to three weeks were required to complete the pollination of an inflorescence. After fruit ripening, hybrid seeds were extracted and sown in the Petri plates for germination. After four weeks, germinated seedlings were transplanted to the portrays, hardened and planted in open field for evaluation. Twenty-five hybrids, yielding more than 1.0 kg fruit weight, were selected for subsequent open field evaluation (latitude 10°55’18.7“N and longitude 76°28’04.6”E). The research station has sandy loam soil, tropical hot and humid weather with a mean annual rainfall of 2788.1 mm. Fruit weight, fruit shape, TSS (°Brix) were evaluated by following randomized block design with two replications, during 2017-2019, along with Kew and check varieties Mauritius and Amritha.

Five suckers of each hybrid, each weighing 500-1000 g, were planted and used for recording 34 parameters as per the Descriptors for pineapple (IBPGR, 1991). The traits plant height, leaves per plant, length, breadth and area of the D leaf, leaf area index, days to attain ideal leaf stage for flowering and days for initiation of flowering (visual) were recorded at flower initiation stage. The traits days for 50 per cent flowering and length of flowering phase were recorded until fruit initiation. The D leaf area (cm^2^) was worked out following the formula suggested by Balakrishnan *et al*. (1978) as (length of ‘D’ leaf x breadth of ‘D’ leaf x 0.725) and Leaf Area Index was worked out following the formula suggested by Watson (1952) as (total leaf area per plant/ total land area occupied per plant).

Fruit traits recorded were length, breadth, girth and weight of the fruits, crown weight, taper ratio, profile of eyes, relative surface of eyes, juice content, peel weight, pulp weight, pulp percentage, days for fruit maturity, crop duration, and shelf life. Fruit length, girth, and breadth were recorded using Vernier calipers whereas weight of freshly harvested fruits including the crown was taken as Fruit weight. Taper ratio of the fruit was the ratio of fruit diameters at ¾ and ¼ height. Profile of eyes (flat, normal, and prominent) and relative surface of the eyes (small, medium, and large) were recorded following the scores given in IBPGR descriptor. Pulp weight was the weight recorded after removing the peel and central core of the fruit whereas pulp percentage was calculated as [pulp weight/ weight of fruit without crown] x 100. Juice content in the fruits (%) was estimated using the formula [weight of juice/ weight of fruit pulp] x 100. Days for fruit maturity in each hybrid was estimated as the mean number of days taken from the opening of first flower to development of three quarters of colour on the fruit when it reaches the standard cultivar size. Crop duration was the number of days taken from date of planting until completion of harvest of all plants of the respective hybrid. Shelf life of the fruits was the number of days from harvest for which the fruits remained marketable when stored at ambient temperature in laboratory conditions. The fruits were rated as not marketable when ≥50 % in a lot were spoiled.

Additionally, fruit TSS, acidity, total sugars, reducing sugars, non-reducing sugars, sugar/ acid ratio, fibre, total carotenoids and ascorbic acid content were estimated by following the standard protocols (AOAC, 2000). All the observations in each hybrid were replicated three times and data analysed using OPSTAT v. 6.0.81.69 (Sheoran et al., 1998).

Average heterosis for any trait in any hybrid was calculated as [(value of the trait in the hybrid – average value of that trait in parents)/ average value of that trait in parents] x 100. Heterobeltiosis for any trait in any hybrid was calculated as [(value of the trait in the hybrid – value of that trait in the better parent)/ value of that trait in the better parent] x 100. Standard heterosis was [(value of the trait in the hybrid – value of that trait in check variety)/value of that trait in check variety] x 100.

To select the best hybrid, a selection criterion (SC) was formulated using the average heterosis values for the economically important fruit traits. SC was calculated as [∑average heterosis (fruit weight with crown, TSS, pulp weight) – ∑average heterosis (crown weight, peel weight, profile of eyes, relative surface of eyes, days to attain ideal leaf stage for flowering, acidity)]. The hybrids identified were also confirmed to meet the basic requirements in fruit weight (≥ 1kg), pulp weight (0.75-0.91 kg), TSS (14.49-15.69 ^○^Brix), days to attain ideal leaf stage for flowering (184.37-185.70 days), crown weight (0.17-0.31 kg), peel weight (0.14-0.16 kg) and acidity (0.81-1.05).

## Results and Discussion

Pineapple is a crop that received comparatively lesser attention towards well designed improvement programmes (Achigan-Dako et al., 2014). Many of the works were restricted to the collection of promising lines and their evaluation for the prioritised locations (Sen, 2001; Hassan et al., 2011; Adje et al., 2019). The inherent difficulties in this crop including the heavy influence of environment and hormonal levels on flowering make the crossing programmes difficult (Van Overbeek and Cruzado, 1948; Bartholomew, 1977; Wang et al., 2007). Further, the breeding programmes have been hindered by the self-incompatibility, poor compatibility among the genotypes and the associated inbreeding depression (Brewbaker and Gorrez, 1967; Bhowmik and Bhagabat, 1975; Coppens d’Eeckenbrugge et al., 1992; Cabral et al., 2003; Sanewski, 2009). Thus, the hybrid development has been comparatively lesser in this crop (Chan, 1991; Chan and Lee, 1999; Cabral et al., 2009; Souza et al., 2009; Hadiati et al., 2011; Viana et al., 2013) and MD-2 (Van de Poel et al., 2009) and FLHORAN41 (Brat et al., 2004) remain as the leading pineapple hybrids in the world.

Even though initial hybridization attempts in pineapple were successful (Kuriakose, 2004), Amritha (Kew x Riply Queen), released by Kerala Agricultural University, India, is the only commercially available hybrid in India. Kew and Mauritius are undoubtedly the leading pineapple cultivars of the world (Kishore et al., 2021) and hence it has been a great idea to cross them and to select for the best hybrid

### Plant and fruit traits of the hybrids

Twenty five Mauritius x Kew hybrids were evaluated under open field conditions, along with their male parent Kew, female parent cum check variety Mauritius and check variety Amritha. For all the characters, there were significant differences among the hybrids. H-35 had the fruit weight of 2.15 kg and H-17 recorded 1.59 kg pulp weight, compared to mid parent values of 1.08 and 0.74 kg, respectively. An increasing in TSS was seen in H-43 (15.75 °Brix). Traits such as days to attain ideal leaf stage for flowering (H-59: 178.14 days), crown weight (H-30: 0.10 kg), peel weight (H-77, H-10 and H-14: 0.08 kg), profile of eyes (H-7: 3.38), relative surface of eyes (H-48: 3.57), and acidity (H-43: 0.81 %), were found to have reduced in the hybrids.

### Heterobeltiosis in hybrids

For plant height and length of D leaf, negative heterobeltiosis was seen on all the hybrids (Table 2). For breadth of D leaf and D leaf area, heterobeltiosis was as high as 52.74 and 39.49 %, respectively in the hybrid H49. In all the hybrids, days taken to physiological maturity and days taken to first flowering had positive heterobeltiosis, showing an extended vegetative growth. For the fruit traits such as length, girth and breadth, the heterobeltiosis was found to be as high as 29.90 (H59), 68.33 (H70) and 28.30 % (H43), respectively. There was a significant heterobeltiosis for crown weight in all the hybrids, which varied from 2.06 (H30) to 429.90 % (H54). Highest heterobeltiosis for pulp weight and pulp percentage were seen in H17. As in case of the vegetative period, all hybrids had positive heterobeltiosis for days for fruit maturity and crop duration, suggesting the extended crop duration in all hybrids. Promising heterobeltiosis of 16.49 % in TSS content (H43), 7.67 % in total sugars (H77), 53.13 % in sugar/acid ratio (H43), 27.37 % in total carotenoids (H70), 60.69 % in ascorbic acid content (H62) and 90.27 % in fruit weight (H35) were also observed among the hybrids. Through diallel crosses, Chan (1991) has developed the hybrids of Queen and Smooth Cayenne, showing heterosis for eight agronomic characters. Sanewski (1998) developed advanced pineapple crosses including PRI hybrid 73-50, with advanced parentage values resulted from the accumulation of desirable genes during earlier crosses and selection processes. Results of the present study confirmed the heterosis previously reported for the commercially important traits in pineapple (d’Eeckenbrugge and Marie, 1998; Marie et al., 1998; Cabral et al., 2000; Brat et al., 2004; Kuriakose, 2004; Fournier et al., 2007; Paull et al., 2017; Viana et al., 2020).

**Table 2.** Heterobeltiosis for plant and fruit traits in Mauritius x Kew hybrids.

| Treatment/<br>Accession | Plant height | No of<br>leaves | Length D<br>leaf | Breadth D<br>leaf | D leaf area | Leaf area<br>Index | Days to<br>attain<br>physiological<br>maturity | Days for<br>flower<br>initiation<br>(Visual) | Days for<br>50 per<br>cent<br>flowering | Flowering<br>phase | Length of<br>the fruit | Girth of the<br>fruit | Breadth of<br>the fruit | Crown<br>weight | Taper<br>ratio | Peel<br>weight | Pulp weight |  |
| --- | --- | --- | --- | --- | --- | --- | --- | --- | --- | --- | --- | --- | --- | --- | --- | --- | --- | --- |
| T 01 | Kew (male parent) |  |  |  |  |  |  |  |  |  |  |  |  |  |  |  |  |  |
| T 02 | Mauritius (check variety cum female parent) |  |  |  |  |  |  |  |  |  |  |  |  |  |  |  |  |  |
| T 03 | Amritha (check variety) |  |  |  |  |  |  |  |  |  |  |  |  |  |  |  |  |  |
| T 04 | H 17 | -14.72 | -2.68 | -12.75 | -13.01 | -24.13 | -26.16 | 4.37 | 18.22 | 3.76 | -9.64 | 6.98 | 20.53 | 23.02 | 258.25 | 8.33 | 6.95 | 110.55 |
| T 05 | H 16 | -13.28 | -5.53 | -10.72 | 13.01 | 0.98 | -4.89 | 4.94 | 17.44 | 3.47 | -13.54 | -31.23 | -3.81 | -6.44 | 76.8 | 5.95 | -32.55 | -40.02 |
| T 06 | H 85 | -14.6 | -5.53 | -7.27 | -6.16 | -13.02 | -18.09 | 3.08 | 8.43 | 9.35 | -8.46 | -21.26 | -7.62 | -4.54 | 26.8 | 9.52 | -30.42 | -45.92 |
| T 07 | H 91 | -15.12 | -3.63 | -13.67 | -25.34 | -35.52 | -37.9 | 4.84 | 15.00 | 12.01 | -2.47 | -8.97 | 11 | 12.14 | 103.09 | 0 | 9.09 | -10.36 |
| T 08 | H 48 | -16.86 | 0.33 | -6.04 | 16.44 | 9.53 | 9.78 | 3.42 | 19.17 | 7.58 | -5.47 | 16.28 | 12.9 | 20.91 | 327.84 | -2.38 | 32.78 | 43.47 |
| T 09 | H 62 | -15.39 | -0.78 | -1.03 | 13.7 | 12.5 | 11.49 | 1.74 | 20.28 | 13.13 | -8.42 | -1.00 | 10.85 | 18.27 | 164.43 | 0 | 66.71 | 6.42 |
| T 10 | H 43 | -13.41 | -7.98 | -8.62 | 36.64 | 25 | 15.16 | 5.25 | 25.77 | 1.19 | -12.98 | -9.63 | 19.65 | 28.30 | 147.42 | 5.95 | -21.12 | 43.26 |
| T 11 | H 66 | -13.37 | -8.22 | -16.21 | 30.48 | 9.08 | 0 | 6.85 | 21.39 | 3.51 | -7.25 | 9.63 | 13.64 | 17.11 | 18.56 | 3.57 | 2.72 | 45.74 |
| T 12 | H 77 | -13.87 | -7.67 | -11.09 | 12.33 | -0.13 | -7.82 | 4.15 | 33.29 | -0.22 | -5.38 | -39.2 | -20.09 | -18.16 | 185.57 | -3.57 | -43.61 | -67.47 |
| T 13 | H 92 | -16.01 | -5.46 | -8.83 | 21.23 | 10.49 | 4.4 | 8.25 | 12.33 | 11.41 | 0.43 | -29.24 | -19.35 | -15.1 | 50.52 | 7.14 | -36.33 | -39.38 |
| T 14 | H 63 | -17.47 | -8.38 | -8.35 | 22.95 | 12.78 | 3.18 | 4.8 | 17.01 | 10 | -9.38 | 8.31 | -5.13 | -2.32 | 8.76 | -10.7 | 4.03 | 4.6 |
| T 15 | H 27 | -17.08 | -8.62 | -8.35 | 21.23 | 11.05 | 1.22 | 1.74 | 23.4 | 3.89 | -6.51 | -31.56 | -12.61 | -6.23 | 259.79 | 1.19 | -28.43 | -48.05 |
| T 16 | H 78 | -16.28 | -4.89 | -12.71 | 15.75 | 0.97 | -4.16 | 4.65 | 9.71 | 8.21 | -3.21 | 0.66 | -1.61 | 3.17 | 256.19 | -3.57 | -15.26 | -20.78 |
| T 17 | H 70 | -17.93 | -1.5 | -6.25 | 24.66 | 16.9 | 15.16 | 8.86 | 14.27 | 4.16 | -9.85 | 1.33 | 68.33 | -0.53 | 162.89 | 10.7 | 25.86 | 73.59 |
| T 18 | H 59 | -13.96 | -3.09 | -10.07 | 12.67 | 1.28 | -1.96 | 7.42 | 18.9 | 12.3 | -8.2 | 29.90 | 8.36 | 10.67 | 53.09 | -1.19 | -6.21 | 41.76 |
| T 19 | H 60 | -15.88 | -4.58 | -5.76 | 18.49 | 11.27 | 6.11 | 7.69 | 25.16 | 7.92 | -6.94 | -34.55 | -23.02 | -13.62 | 189.69 | 7.14 | -37.18 | -69.78 |
| T 20 | H 49 | -14.26 | -7.91 | -8.67 | 52.74 | 39.49 | 28.36 | 5.05 | 10.47 | 2.33 | -2.78 | -24.92 | -7.62 | -2.53 | 89.69 | -2.38 | -28.35 | -39.76 |
| T 21 | H 54 | -14.66 | -7.91 | -8.14 | 16.44 | 6.99 | -1.71 | 3.58 | 19.45 | 10.22 | -12.89 | -8.31 | 16.42 | -9.19 | 429.90 | 5.95 | 0.07 | -0.99 |
| T 22 | H 10 | -15.7 | -4.28 | -16.37 | 7.88 | -9.53 | -13.69 | 4.28 | 18.07 | 5.62 | 0.22 | -31.56 | -12.17 | -7.39 | 180.93 | 5.95 | -41.24 | -52.61 |
| T 23 | H 15 | -15.33 | -4.35 | -4.84 | 25.68 | 19.55 | 14.18 | 4.37 | 23.68 | 15.19 | -5.08 | -28.57 | -11.41 | -9.5 | 155.15 | 7.14 | -30.98 | -55.95 |
| T 24 | H 30 | -15.39 | -8.08 | -8.83 | 21.23 | 10.52 | 1.71 | 7.81 | 4.18 | 8.28 | -5.64 | -30.56 | -22.29 | -36.85 | 2.06 | 2.38 | -11.81 | -59.64 |
| T 25 | H 14 | -16.25 | -8.31 | -7.59 | 0.68 | -7.22 | -14.91 | 1.98 | 23.98 | 12.84 | 2.43 | -27.91 | 2.2 | 3.91 | 220.1 | 7.14 | -40.89 | -27.57 |
| T 26 | H 7 | -17.47 | -3.16 | -8.72 | 13.36 | 3.5 | 0 | 6.23 | 12.41 | 14.7 | -6.73 | -16.61 | 0.44 | 1.58 | 123.2 | 5.95 | -12.7 | -23.82 |
| T 27 | H 35 | -15.94 | -0.86 | -10.72 | 22.26 | 9.09 | 8.07 | 3.89 | 14.59 | 11.81 | -9.64 | -29.9 | 26.69 | 1.58 | 242.67 | 5.95 | 27.42 | 71.21 |
| T 28 | H 19 | -13.37 | -6.79 | -5.49 | 46.23 | 38.17 | 28.61 | 4.21 | 22.92 | 13.6 | -10.24 | -26.25 | -15.4 | -5.49 | 91.72 | 7.14 | -17.24 | -20.64 |

| Treatment/<br>Accession | Pulp<br>percentage | Days for<br>fruit<br>maturity | Crop<br>duration | Shelf<br>life | Juice<br>percen<br>tage | TSS | Acidity | Total<br>sugars | Reduc<br>ing<br>sugars | Non-reducing<br>sugars | Sugar/<br>acid<br>ratio | Fibre | Total<br>carotenoids | Ascorbic<br>acid | Fruit<br>weight | Eye<br>profile | Eye<br>relative<br>surface |  |
| --- | --- | --- | --- | --- | --- | --- | --- | --- | --- | --- | --- | --- | --- | --- | --- | --- | --- | --- |
| T 01 | Kew (male parent) |  |  |  |  |  |  |  |  |  |  |  |  |  |  |  |  |  |
| T 02 | Mauritius (check variety cum female parent) |  |  |  |  |  |  |  |  |  |  |  |  |  |  |  |  |  |
| T 03 | Amritha (check variety) |  |  |  |  |  |  |  |  |  |  |  |  |  |  |  |  |  |
| T 04 | H 17 | 9.19 | 6.57 | 6.79 | -5.54 | -13.56 | 7.47 | -3.41 | -2.17 | -10.98 | -90.13 | -82.88 | 0.67 | 7.98 | -23.75 | 87.61 | -6.25 | -12.30 |
| T 05 | H 16 | -6.06 | 3.87 | 5.93 | -2.9 | -0.47 | -8.14 | 4.55 | -14.76 | -35.37 | -22.99 | -5.01 | -19.09 | -0.46 | -50.92 | -32.74 | 10.50 | 11.86 |
| T 06 | H 85 | -10.67 | 3.57 | 3.87 | -3.27 | 5.98 | -1.48 | 1.14 | -10.09 | 0.49 | -32.15 | -19.55 | -15.58 | -2.79 | -65.97 | -39.82 | -10.50 | -1.12 |
| T 07 | H 91 | -10.59 | 2.43 | 5.04 | 0.63 | -3.71 | 10.28 | 3.41 | -2.84 | -7.56 | -20.04 | -17.33 | -6.62 | 5.19 | -41.97 | -0.88 | 59.25 | 9.62 |
| T 08 | H 48 | -2.76 | 3.49 | 5.22 | -7.05 | -6.69 | -2.29 | -5.68 | 0.33 | -36.34 | -4.07 | -20.04 | 2.93 | 4.85 | 21.38 | 56.64 | 38.00 | 9.84 |
| T 09 | H 62 | -14.33 | 7.42 | 6.05 | 13.35 | -2.43 | 0.52 | 19.32 | -2.75 | -24.15 | -13.02 | -15.87 | -0.79 | -5.86 | 60.69 | 23.89 | -6.75 | 35.79 |
| T 10 | H 43 | 4.35 | 1.69 | 6.17 | -13.85 | -9.87 | 16.49 | -7.95 | -7.67 | -13.9 | -23.19 | 53.10 | 1.42 | 9.19 | -19.26 | 32.74 | -10.00 | 11.19 |
| T 11 | H 66 | 0.22 | 8.09 | 8.97 | 0.76 | -1.94 | 3.33 | -2.27 | 6.84 | -45.37 | 7.53 | -20.25 | 6.53 | 22.32 | 7.66 | 28.32 | -4.25 | -20.13 |
| T 12 | H 77 | -26.2 | 3.02 | 6.99 | 0.76 | -2.47 | 1.26 | 0 | 7.67 | -41.22 | 6.82 | -13.29 | 0.39 | -12.52 | 2.36 | -39.82 | 15.00 | 11.86 |
| T 13 | H 92 | -11.59 | 6.43 | 7.99 | 2.9 | -5.25 | 0.81 | 13.64 | -22.6 | -12.93 | -41.81 | -30.76 | -3.87 | 7.88 | 41.69 | -30.97 | 20.00 | 11.86 |
| T 14 | H 63 | 0.6 | 7.13 | 7.09 | -7.05 | 1.47 | -10.06 | 5.68 | -16.85 | -36.83 | -24.92 | -20.25 | 1.57 | 8.48 | 40.76 | -6.19 | 58.25 | -10.51 |
| T 15 | H 27 | -9.38 | 8.22 | 6.71 | 0.38 | -2.1 | -3.33 | 1.14 | 2.84 | -31.46 | -3.15 | -25.12 | -0.39 | -6.46 | 16.89 | -22.12 | 28.25 | 0.00 |
| T 16 | H 78 | -5.62 | 8.94 | 6.9 | -0.5 | -2.91 | -2.07 | 14.77 | -9.01 | -16.59 | -23.8 | -20.88 | -0.73 | 2.63 | -16.36 | -0.88 | 25.00 | 11.86 |
| T 17 | H 70 | 1.51 | 3.64 | 7.43 | -2.9 | -5.52 | -0.89 | -4.55 | -11.18 | -27.32 | -21.97 | -30.83 | 3.63 | 27.37 | 5.8 | 61.95 | -3.50 | -3.58 |
| T 18 | H 59 | 0.87 | 10.83 | 10.04 | -5.29 | -7.97 | 5.7 | -1.14 | -54.96 | -18.54 | -79.04 | -77.52 | 1.36 | 10.5 | -30.87 | 27.43 | -2.25 | -12.53 |
| T 19 | H 60 | -17.25 | 6.33 | 9.13 | -2.39 | -4.69 | 13.02 | -1.14 | -24.94 | -19.27 | -42.12 | -6.26 | 0.54 | 11.31 | -36.67 | -45.13 | 25.00 | 11.86 |
| T 20 | H 49 | -11.18 | 13.91 | 9.13 | 13.35 | -1.73 | 9.39 | 9.09 | -16.1 | -9.27 | -35.5 | -13.92 | -11.04 | -11.13 | -18.48 | -28.32 | 38.25 | 0.00 |
| T 21 | H 54 | -5.22 | 5.07 | 5.95 | 13.35 | -5.6 | 2.96 | 2.27 | -19.77 | -28.05 | -32.15 | -29.71 | 5.69 | 10.61 | 26.13 | 30.09 | 38.25 | 0.00 |
| T 22 | H 10 | -8.34 | 10.03 | 8.08 | -14.99 | -3.25 | 8.36 | 3.41 | -11.93 | -30.24 | -21.67 | -33.12 | -1.33 | 6.06 | 44.57 | -33.63 | -1.75 | 7.38 |
| T 23 | H 15 | -11.62 | 5.59 | 7.02 | -2.39 | -0.21 | 1.26 | 0 | 7.67 | -41.22 | 6.82 | -13.29 | 0.39 | -12.52 | 2.36 | -37.17 | 25.00 | 11.86 |
| T 24 | H 30 | -22.04 | 8.46 | 7.65 | 7.05 | 0.66 | -3.25 | 12.5 | -1.5 | -29.27 | -9.46 | -6.26 | 0.54 | 11.31 | 3.3 | -49.56 | 6.75 | 16.33 |
| T 25 | H 14 | -8.57 | 10.87 | 7.94 | -3.78 | -7.03 | 6.8 | 10.23 | -3.59 | -26.34 | -13.12 | 1.95 | 3.93 | -15.76 | 29.02 | -7.96 | 41.75 | -3.13 |
| T 26 | H 7 | -12.25 | 10.3 | 8.51 | -11.84 | -4.07 | 2.88 | 17.05 | -10.68 | -33.66 | -18.82 | -13.29 | 0.33 | -18.48 | -1.85 | -10.62 | -15.50 | 6.26 |
| T 27 | H 35 | -13.63 | 5.63 | 5.78 | -7.56 | -5.3 | 11.46 | -2.27 | 6.59 | -53.17 | 10.58 | 16.42 | 0.27 | -6.57 | 13.2 | 90.27 | -1.75 | 3.13 |
| T 28 | H 19 | -4.17 | 6.01 | 7.02 | -1.26 | 4.66 | 8.51 | 1.14 | 3 | -46.1 | 3.26 | 3.62 | 2.9 | 12.12 | 31.39 | -15.93 | 1.75 | -5.82 |

### Standard heterosis over Mauritius

Compared to Mauritius, all the hybrids had shown negative heterosis for plant height, number of leaves and length of the D leaf (Table 3). Hybrids H59 (29.90 %), H70 (79.94 %) and H43 (46.21 %) had shown maximum positive standard heterosis for fruit length, girth and breadth, respectively. Maximum heterosis for pulp weight, pulp percentage, TSS and sugar/acid ratio were shown by H17 (117.99, 14.19, 16.49 and 53.10 %, respectively). H70 had highest heterosis (42.49 %) for highest fruit carotenoid content whereas H62 had highest for ascorbic acid content (60.69 %).

**Table 3.**
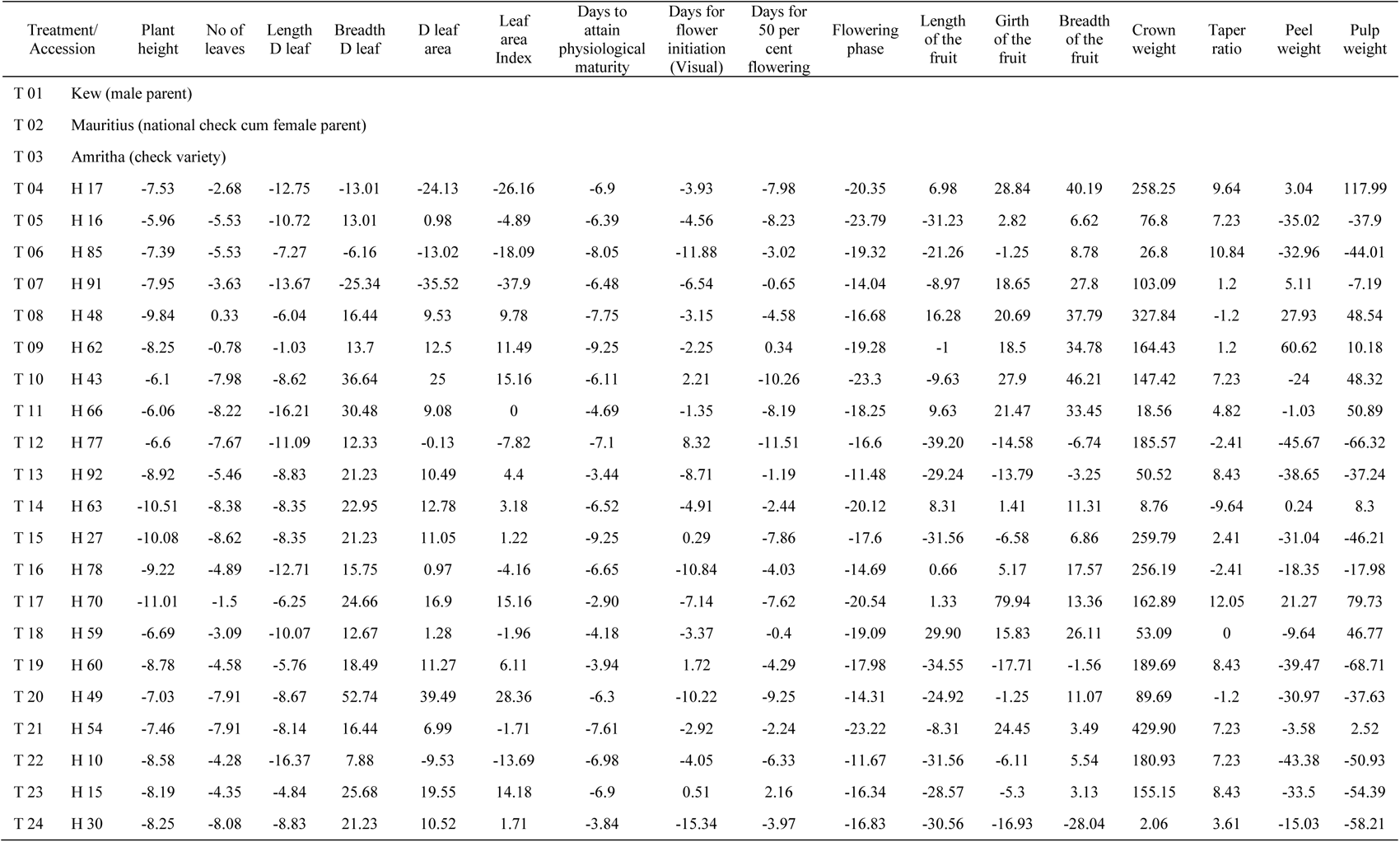
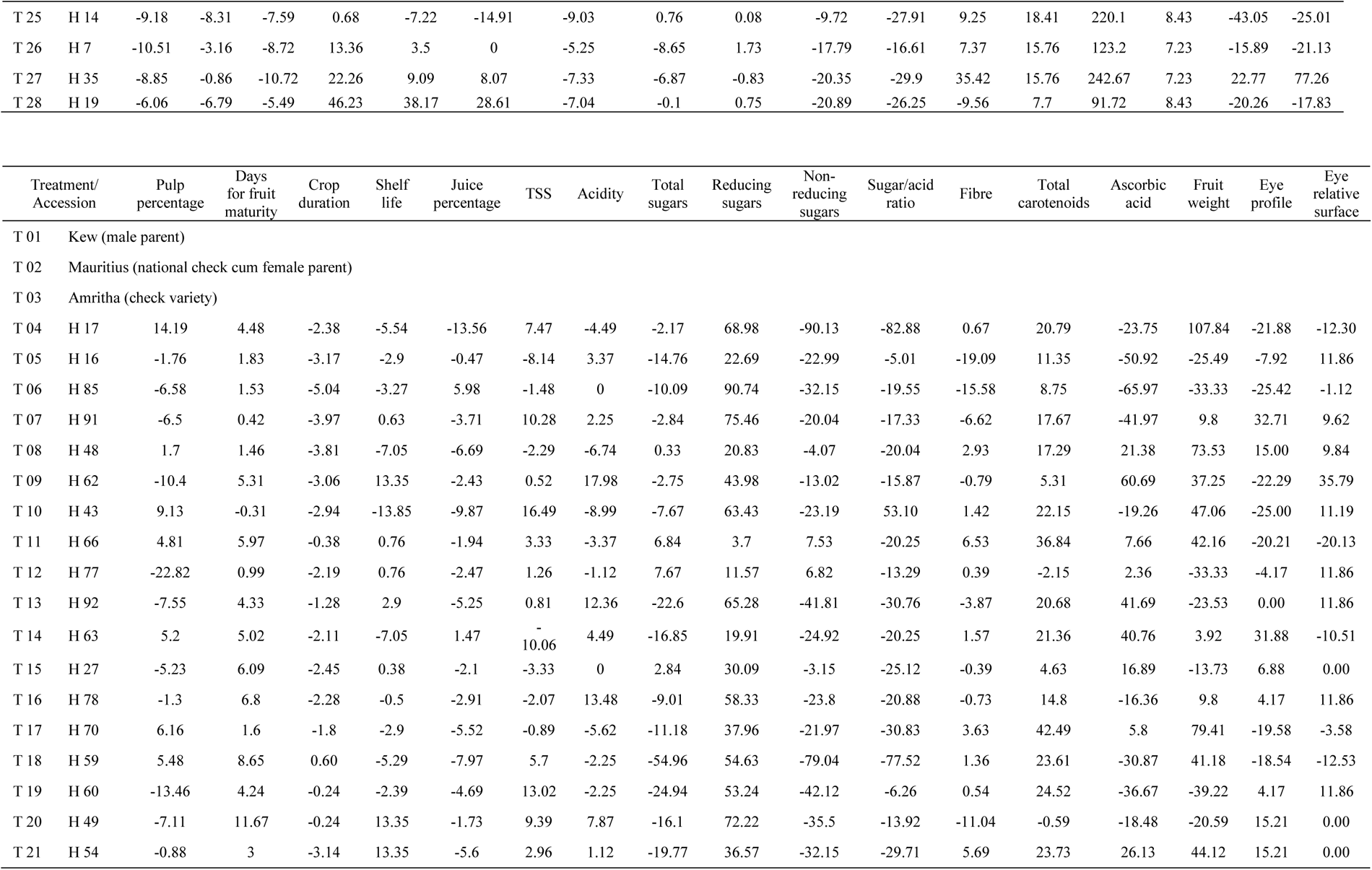
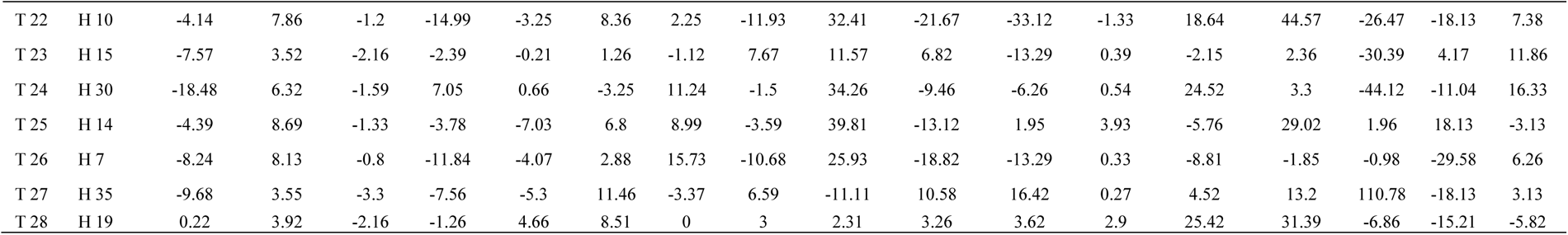
Standard heterosis for plant and fruit traits in Mauritius x Kew hybrids over the check variety cum female parent Mauritius.

### Standard heterosis over Amritha

Over the check parent Amritha, the hybrids had high standard heterosis for D leaf area and leaf area index. Maximum heterosis for these traits were seen in H49 (82.45 %) and H19 (74.75 %), respectively (Table 4). Days taken for flower initiation and 50 per cent flowering had slightly lower heterosis over Amritha. The fruit traits such as length, girth and breadth had shown positive heterosis with maximum in H59 (67.81 %), H70 (91.33 %) and H43 (45.68 %). The higher standard heterosis for crown weight in all the hybrids is viewed as a disadvantage. Heterosis for pulp weight and pulp percentage were highest in H17 (293.34 and 22.75 %, respectively). Almost all the hybrids have shown better shelf life but the reduced TSS content in the hybrids was another disadvantage over the check variety. Heterosis for sugar/acid ratio, carotenoid content, ascorbic acid content and fruit weight were highest in H43 (72.01 %), H70 (21.60 %), H62 (107.85 %) and H35 (230.77 %), respectively.

**Table 4.** Standard heterosis for plant and fruit traits in Mauritius x Kew hybrids over the check variety Amritha.

| Treatment/<br>Accession | Plant<br>height | No of<br>leaves | Length<br>D leaf | Breadth<br>D leaf | D leaf<br>area | Leaf<br>area<br>Index | Days to<br>attain<br>physiological<br>maturity | Days for<br>flower<br>initiation<br>(Visual) | Days for<br>50 per<br>cent<br>flowering | Flowering<br>phase | Length<br>of the<br>fruit | Girth<br>of the<br>fruit | Breadth<br>of the<br>fruit | Crown<br>weight | Taper<br>ratio | Peel<br>weight | Pulp<br>weight |  |
| --- | --- | --- | --- | --- | --- | --- | --- | --- | --- | --- | --- | --- | --- | --- | --- | --- | --- | --- |
| T 01 | Kew (male parent) |  |  |  |  |  |  |  |  |  |  |  |  |  |  |  |  |  |
| T 02 | Mauritius (national check cum female parent) |  |  |  |  |  |  |  |  |  |  |  |  |  |  |  |  |  |
| T 03 | Amritha (check variety) |  |  |  |  |  |  |  |  |  |  |  |  |  |  |  |  |  |
| T 04 | H 17 | 1.76 | 0.81 | 3.05 | -3.79 | -0.76 | 0.33 | 1.21 | -4.90 | -4.96 | -16.62 | 38.20 | 37.00 | 39.69 | 249.25 | 7.06 | 21.83 | 293.34 |
| T 05 | H 16 | 3.48 | -2.14 | 5.45 | 25.00 | 32.09 | 29.24 | 1.76 | -5.53 | -5.23 | -20.22 | -11.16 | 9.33 | 6.24 | 72.36 | 4.71 | -23.17 | 12.06 |
| T 06 | H 85 | 1.90 | -2.14 | 9.53 | 3.79 | 13.77 | 11.30 | -0.04 | -12.77 | 0.16 | -15.54 | 1.72 | 5.00 | 8.39 | 23.62 | 8.24 | -20.74 | 1.03 |
| T 07 | H 91 | 1.29 | -0.17 | 1.96 | -17.42 | -15.66 | -15.61 | 1.66 | -7.49 | 2.60 | -10.01 | 17.60 | 26.17 | 27.34 | 97.99 | -1.18 | 24.27 | 67.47 |
| T 08 | H 48 | -0.79 | 3.94 | 10.98 | 28.79 | 43.26 | 49.17 | 0.28 | -4.13 | -1.45 | -12.78 | 50.21 | 28.33 | 37.29 | 317.09 | -3.53 | 51.25 | 168.03 |
| T 09 | H 62 | 0.96 | 2.78 | 16.90 | 25.76 | 47.15 | 51.50 | -1.35 | -3.24 | 3.63 | -15.50 | 27.90 | 26.00 | 34.29 | 157.79 | -1.18 | 89.91 | 98.81 |
| T 10 | H 43 | 3.33 | -4.68 | 7.93 | 51.14 | 63.50 | 56.48 | 2.07 | 1.17 | -7.32 | -19.70 | 16.74 | 36.00 | 45.68 | 141.21 | 4.71 | -10.15 | 167.63 |
| T 11 | H 66 | 3.37 | -4.92 | -1.03 | 44.32 | 42.67 | 35.88 | 3.62 | -2.35 | -5.18 | -14.42 | 41.63 | 29.17 | 32.97 | 15.58 | 2.35 | 17.02 | 172.26 |
| T 12 | H 77 | 2.78 | -4.36 | 5.02 | 24.24 | 30.63 | 25.25 | 1.00 | 7.23 | -8.61 | -12.70 | -21.46 | -9.17 | -7.07 | 178.39 | -4.71 | -35.77 | -39.23 |
| T 13 | H 92 | 0.22 | -2.07 | 7.68 | 34.09 | 44.52 | 41.86 | 4.97 | -9.64 | 2.05 | -7.33 | -8.58 | -8.33 | -3.60 | 46.73 | 5.88 | -27.47 | 13.24 |
| T 14 | H 63 | -1.53 | -5.09 | 8.26 | 35.98 | 47.52 | 40.20 | 1.62 | -5.87 | 0.76 | -16.38 | 39.91 | 7.83 | 10.91 | 6.03 | -11.76 | 18.51 | 95.41 |
| T 15 | H 27 | -1.05 | -5.34 | 8.26 | 34.09 | 45.25 | 37.54 | -1.35 | -0.73 | -4.84 | -13.74 | -11.59 | -0.67 | 6.47 | 250.75 | 0.00 | -18.47 | -2.95 |
| T 16 | H 78 | -0.10 | -1.48 | 3.11 | 28.03 | 32.07 | 30.23 | 1.48 | -11.74 | -0.88 | -10.69 | 30.04 | 11.83 | 17.15 | 247.24 | -4.71 | -3.47 | 48.00 |
| T 17 | H 70 | -2.07 | 2.04 | 10.74 | 37.88 | 52.91 | 56.48 | 5.56 | -8.08 | -4.59 | -16.82 | 30.90 | 91.33 | 12.95 | 156.28 | 9.41 | 43.38 | 224.30 |
| T 18 | H 59 | 2.67 | 0.39 | 6.22 | 24.62 | 32.48 | 33.22 | 4.17 | -4.35 | 2.87 | -15.30 | 67.81 | 23.17 | 25.66 | 49.25 | -2.35 | 6.84 | 164.83 |
| T 19 | H 60 | 0.37 | -1.16 | 11.31 | 31.06 | 45.54 | 44.19 | 4.43 | 0.69 | -1.15 | -14.14 | -15.45 | -12.50 | -1.92 | 182.41 | 5.88 | -28.43 | -43.55 |
| T 20 | H 49 | 2.30 | -4.60 | 7.88 | 68.94 | 82.45 | 74.42 | 1.87 | -11.13 | -6.27 | -10.29 | -3.00 | 5.00 | 10.67 | 84.92 | -3.53 | -18.38 | 12.53 |
| T 21 | H 54 | 1.83 | -4.60 | 8.50 | 28.79 | 39.94 | 33.55 | 0.44 | -3.91 | 0.96 | -19.62 | 18.45 | 32.33 | 3.12 | 416.58 | 4.71 | 13.99 | 84.98 |
| T 22 | H 10 | 0.59 | -0.84 | -1.22 | 19.32 | 18.34 | 17.28 | 1.12 | -5.02 | -3.26 | -7.53 | -11.59 | -0.17 | 5.16 | 173.87 | 4.71 | -33.06 | -11.46 |
| T 23 | H 15 | 1.03 | -0.91 | 12.40 | 39.02 | 56.37 | 55.15 | 1.21 | -0.51 | 5.51 | -12.41 | -7.73 | 0.70 | 2.76 | 148.74 | 5.88 | -21.37 | -17.70 |
| T 24 | H 30 | 0.96 | -4.77 | 7.68 | 34.09 | 44.56 | 38.21 | 4.54 | -16.19 | -0.82 | -12.94 | -10.30 | -11.67 | -28.30 | -0.50 | 1.18 | 0.46 | -24.60 |
| T 25 | H 14 | -0.07 | -5.02 | 9.15 | 11.36 | 21.35 | 15.61 | -1.11 | -0.26 | 3.36 | -5.49 | -6.87 | 16.17 | 17.99 | 212.06 | 5.88 | -32.66 | 35.31 |
| T 26 | H 7 | -1.53 | 0.32 | 7.82 | 25.38 | 35.37 | 35.88 | 3.01 | -9.57 | 5.06 | -13.94 | 7.73 | 14.17 | 15.35 | 117.59 | 4.71 | -0.55 | 42.32 |
| T 27 | H 35 | 0.30 | 2.71 | 5.45 | 35.23 | 42.68 | 46.84 | 0.75 | -7.81 | 2.42 | -16.62 | -9.44 | 44.00 | 15.35 | 234.06 | 4.71 | 45.15 | 219.84 |
| T 28 | H 19 | 3.37 | -3.44 | 11.63 | 61.74 | 80.73 | 74.75 | 1.06 | -1.11 | 4.06 | -17.18 | -4.72 | -3.83 | 7.31 | 86.90 | 5.88 | -5.73 | 48.26 |

| Treatment/<br>Accession | Pulp<br>percentage | Days<br>for fruit<br>maturity | Crop<br>duration | Shelf<br>life | Juice<br>percentage | TSS | Acidity | Total<br>sugars | Reducing<br>sugars | Non-reducing<br>sugars | Sugar/acid<br>ratio | Fibre | Total<br>carotenoids | Ascorbic<br>acid | Fruit<br>weight | Eye<br>profile | Eye<br>relative<br>surface |  |
| --- | --- | --- | --- | --- | --- | --- | --- | --- | --- | --- | --- | --- | --- | --- | --- | --- | --- | --- |
| T 01 | Kew (male parent) |  |  |  |  |  |  |  |  |  |  |  |  |  |  |  |  |  |
| T 02 | Mauritius (national check cum female parent) |  |  |  |  |  |  |  |  |  |  |  |  |  |  |  |  |  |
| T 03 | Amritha (check variety) |  |  |  |  |  |  |  |  |  |  |  |  |  |  |  |  |  |
| T 04 | H 17 | 22.75 | 4.40 | 1.60 | 3.45 | -11.68 | -13.61 | -1.16 | 8.01 | 11.28 | -87.20 | -80.77 | -4.12 | 3.08 | -1.38 | 226.15 | -25.00 | -21.60 |
| T 05 | H 16 | 5.61 | 1.75 | 0.78 | 6.34 | 1.71 | -26.16 | 6.98 | -5.89 | -19.21 | -0.13 | 6.72 | -22.93 | -4.97 | -36.51 | 16.92 | -11.60 | 0.00 |
| T 06 | H 85 | 0.42 | 1.46 | -1.17 | 5.93 | 8.30 | -20.81 | 3.49 | -0.74 | 25.61 | -12.01 | -9.62 | -19.59 | -7.19 | -55.98 | 4.62 | -28.40 | -11.60 |
| T 07 | H 91 | 0.51 | 0.35 | -0.06 | 10.21 | -1.60 | -11.36 | 5.81 | 7.27 | 15.55 | 3.69 | -7.11 | -11.06 | 0.42 | -24.94 | 72.31 | 27.40 | -2.00 |
| T 08 | H 48 | 9.32 | 1.38 | 0.11 | 1.79 | -4.65 | -21.46 | -3.49 | 10.77 | -20.43 | 24.41 | -10.16 | -1.96 | 0.10 | 57.00 | 172.31 | 10.40 | -1.80 |
| T 09 | H 62 | -3.69 | 5.23 | 0.90 | 24.14 | -0.30 | -19.20 | 22.09 | 7.37 | -5.18 | 12.80 | -5.47 | -5.50 | -10.13 | 107.85 | 115.38 | -25.40 | 21.40 |
| T 10 | H 43 | 17.31 | -0.38 | 1.02 | -5.66 | -7.90 | -6.36 | -5.81 | 1.93 | 7.62 | -0.40 | 72.01 | -3.40 | 4.24 | 4.43 | 130.77 | -28.00 | -0.60 |
| T 11 | H 66 | 12.66 | 5.89 | 3.68 | 10.34 | 0.20 | -16.94 | 0.00 | 17.96 | -31.71 | 39.45 | -10.40 | 1.47 | 16.78 | 39.25 | 123.08 | -23.40 | -28.60 |
| T 12 | H 77 | -17.04 | 0.92 | 1.80 | 10.34 | -0.34 | -18.61 | 2.33 | 18.88 | -26.52 | 38.52 | -2.58 | -4.38 | -16.49 | 32.40 | 4.62 | -8.00 | 0.00 |
| T 13 | H 92 | -0.62 | 4.26 | 2.75 | 12.69 | -3.18 | -18.97 | 16.28 | -14.55 | 8.84 | -24.54 | -22.20 | -8.44 | 2.99 | 83.27 | 20.00 | -4.00 | 0.00 |
| T 14 | H 63 | 13.09 | 4.95 | 1.89 | 1.79 | 3.69 | -27.71 | 8.14 | -8.20 | -21.04 | -2.64 | -10.40 | -3.26 | 3.57 | 82.07 | 63.08 | 26.60 | -20.00 |
| T 15 | H 27 | 1.88 | 6.01 | 1.53 | 9.93 | 0.03 | -22.29 | 3.49 | 13.54 | -14.33 | 25.59 | -15.87 | -5.13 | -10.70 | 51.19 | 35.38 | 2.60 | -10.60 |
| T 16 | H 78 | 6.10 | 6.72 | 1.71 | 8.97 | -0.79 | -21.28 | 17.44 | 0.46 | 4.27 | -1.19 | -11.10 | -5.45 | -2.03 | 8.19 | 72.31 | 0.00 | 0.00 |
| T 17 | H 70 | 14.12 | 1.53 | 2.21 | 6.34 | -3.46 | -20.33 | -2.33 | -1.93 | -9.15 | 1.19 | -22.28 | -1.30 | 21.60 | 36.85 | 181.54 | -22.80 | -13.80 |
| T 18 | H 59 | 13.39 | 8.57 | 4.70 | 3.72 | -5.96 | -15.04 | 1.16 | -50.28 | 1.83 | -72.82 | -74.75 | -3.46 | 5.49 | -10.58 | 121.54 | -21.80 | -21.80 |
| T 19 | H 60 | -6.98 | 4.17 | 3.83 | 6.90 | -2.61 | -9.16 | 1.16 | -17.13 | 0.91 | -24.93 | 5.32 | -4.24 | 6.27 | -18.09 | -4.62 | 0.00 | 0.00 |
| T 20 | H 49 | -0.15 | 11.59 | 3.83 | 24.14 | 0.42 | -12.07 | 11.63 | -7.37 | 13.41 | -16.36 | -3.28 | -15.27 | -15.16 | 5.45 | 24.62 | 10.60 | -10.60 |
| T 21 | H 54 | 6.55 | 2.93 | 0.81 | 24.14 | -3.54 | -17.24 | 4.65 | -11.42 | -10.06 | -12.01 | -21.03 | 0.66 | 5.59 | 63.15 | 126.15 | 10.60 | -10.60 |
| T 22 | H 10 | 3.04 | 7.79 | 2.83 | -6.90 | -1.14 | -12.90 | 5.81 | -2.76 | -12.80 | 1.58 | -24.86 | -6.02 | 1.25 | 87.00 | 15.38 | -21.40 | -4.00 |
| T 23 | H 15 | -0.64 | 3.44 | 1.83 | 6.90 | 1.97 | -18.61 | 2.33 | 18.88 | -26.52 | 38.52 | -2.58 | -4.38 | -16.49 | 32.40 | 9.23 | 0.00 | 0.00 |
| T 24 | H 30 | -12.36 | 6.25 | 2.43 | 17.24 | 2.86 | -22.24 | 15.12 | 8.75 | -11.59 | 17.41 | 5.32 | -4.24 | 6.27 | 33.62 | -12.31 | -14.60 | 4.00 |
| T 25 | H 14 | 2.78 | 8.61 | 2.70 | 5.38 | -5.00 | -14.15 | 12.79 | 6.45 | -7.93 | 12.66 | 14.54 | -1.01 | -19.58 | 66.88 | 60.00 | 13.40 | -13.40 |
| T 26 | H 7 | -1.36 | 8.05 | 3.24 | -3.45 | -1.97 | -17.30 | 19.77 | -1.38 | -17.07 | 5.28 | -2.58 | -4.44 | -22.18 | 26.95 | 55.38 | -32.40 | -5.00 |
| T 27 | H 35 | -2.91 | 3.48 | 0.64 | 1.24 | -3.23 | -10.40 | 0.00 | 17.68 | -41.46 | 43.40 | 30.81 | -4.49 | -10.80 | 46.42 | 230.77 | -21.40 | -7.80 |
| T 28 | H 19 | 7.73 | 3.85 | 1.83 | 8.14 | 6.94 | -12.78 | 3.49 | 13.72 | -32.62 | 33.91 | 16.42 | -1.99 | 7.04 | 69.95 | 46.15 | -18.60 | -15.80 |

### Average heterosis in hybrids

All the hybrids have shown negative average heterosis for plant height but the leaf area index was found to have improved with the highest in H19 (40.83 %) (Table 5). Days taken for flower initiation and 50 per cent flowering were almost similar to the parents, with the maximum heterosis of 19.52 (H77) and 1.79 % (H30), respectively. For fruit length, girth and breadth, there were mixed responses from the hybrids however, the maximum heterosis for these traits were shown by H59 (40.65 %), H70 (73.94 %) and H43 (36.67 %), respectively. Positive heterosis for crown weight shown by most of the hybrids was an unfavourable factor in the selection of hybrids. Pulp weight and pulp percentage had highest heterosis in H17 (114.21 and 11.64 %, respectively). Crop duration in all the hybrids was nearly same as in the parents, with the heterosis values ranging between -0.79 (H85) and 5.11 % (H59). In most of the hybrids, TSS, acidity and sugar/acid ratio were improved with the highest heterosis of 21.2 (H43), 18.64 (H62) and 60.76 % (H43), respectively. Total carotenoids, ascorbic acid content and fruit weight were also improved in most of the hybrids with the maximum heterosis of 34.51 (H70), 90.92 (H62) and 100.00 % (H35), respectively.

**Table 5.**
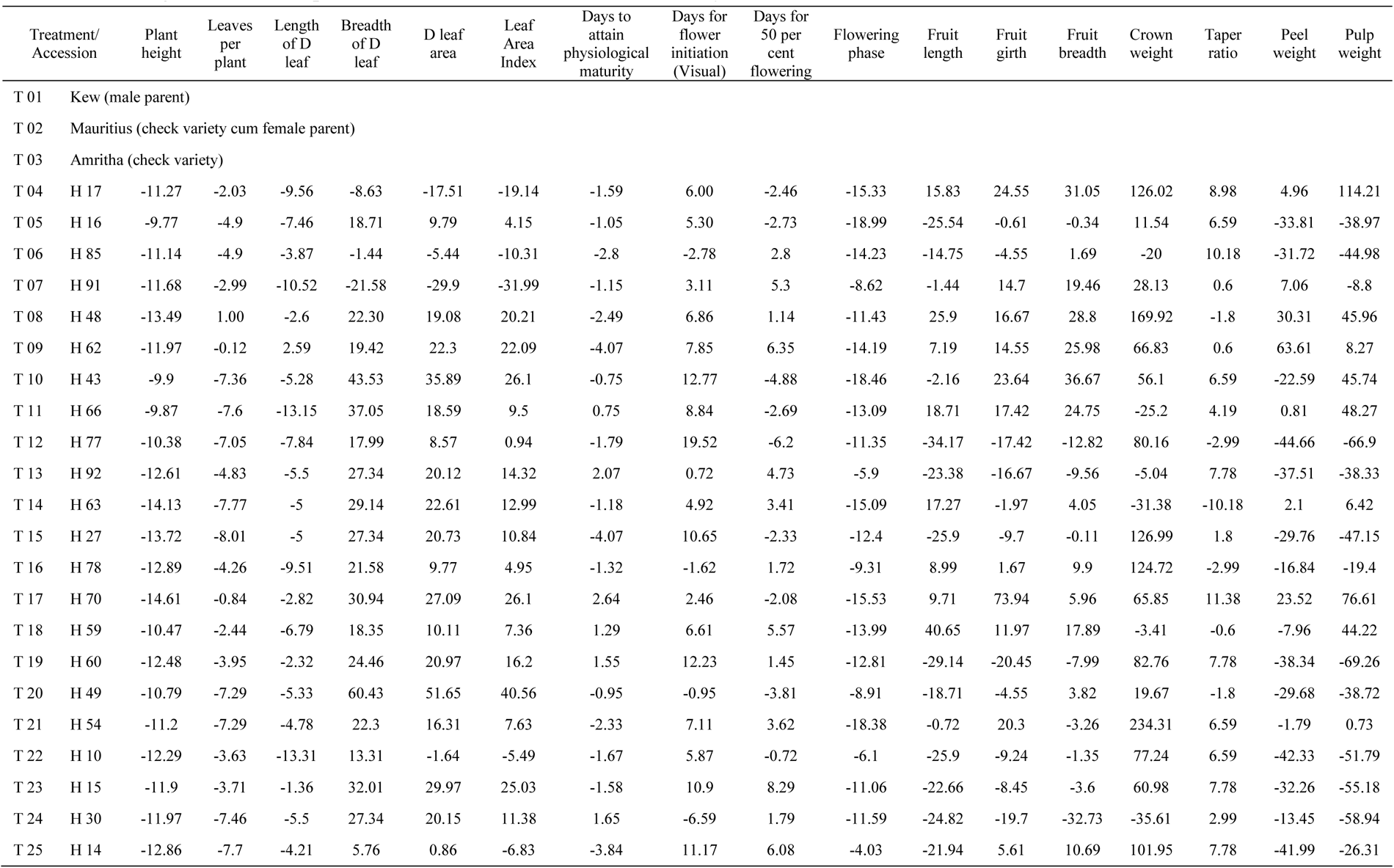
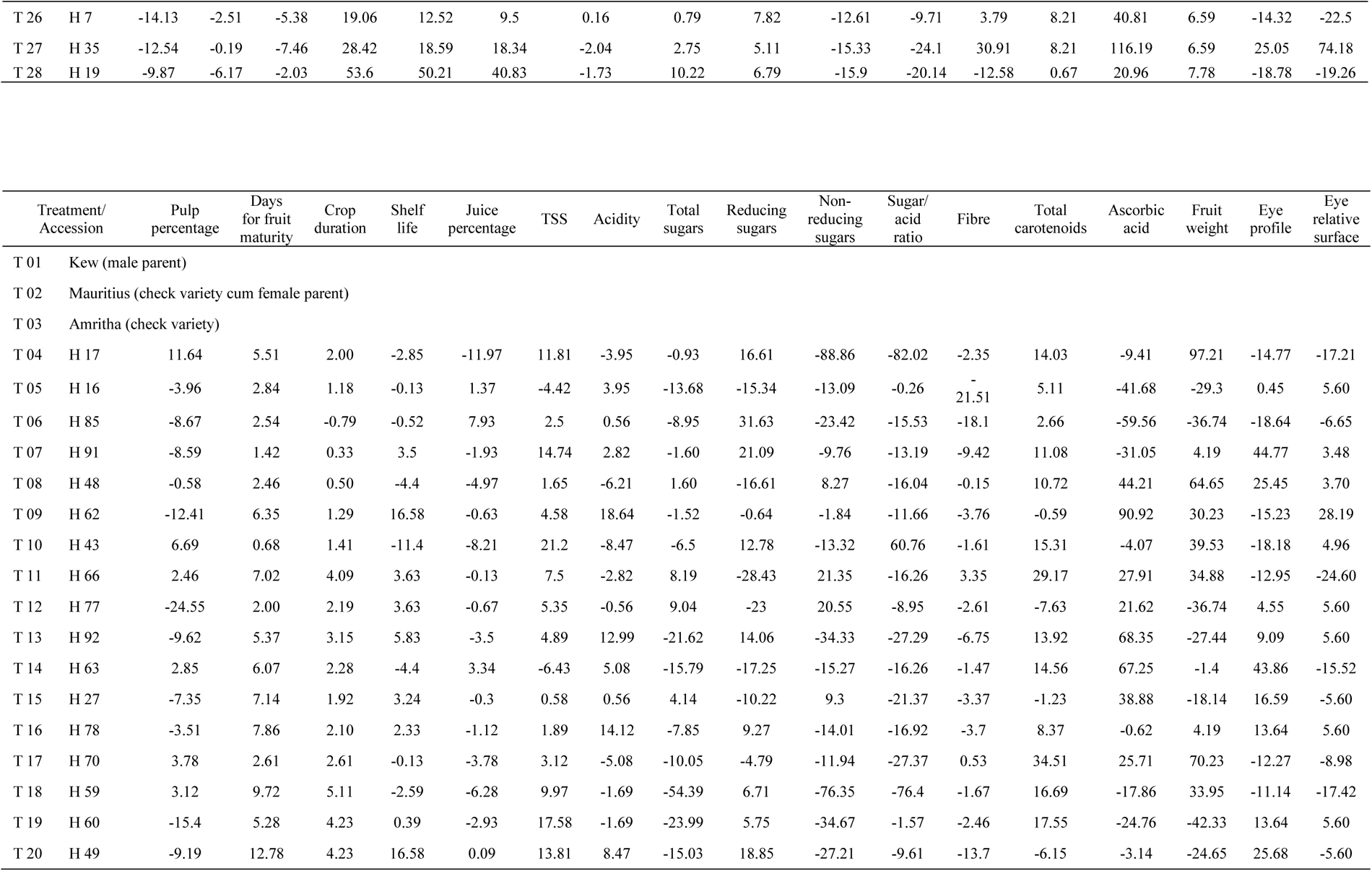
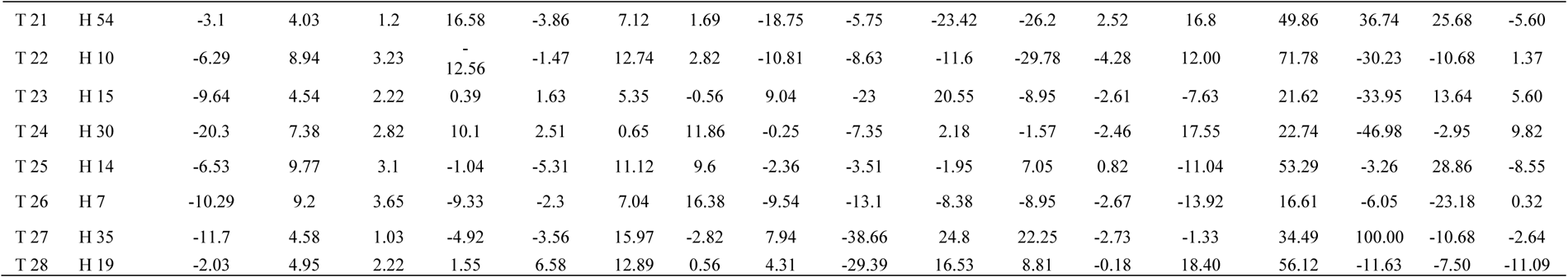
Average heterosis for plant and fruit traits in Mauritius x Kew hybrids.

### Selection of hybrids

Improvement in fruit weight and quality is the leading breeding objective in pineapple (Kuriakose, 2004). The selection index provides appropriate weightage to the phenotypic values of multiple characters in selection process (Fisher, 1936; de Souza et al., 2000). Often, selection criteria, multiple regression relations or indices are more efficient than individual selection based on individual traits (Moreira et al., (2019).

Thus, in this study also, selection criterion was formulated based on multiple desirable and undesirable fruit and plant traits. Using the selection criterion, values were calculated for all the hybrids (Table 6). The highest values were observed in H66 (154.66), H17 (129.77), H59 (128.47), H43 (95.40), H70 (84.28) and H35 (67.09). All these hybrids were found to meet the requirements in fruit weight (≥1.0 kg), pulp weight (≥0.75 kg), TSS (≥14.49 ^○^Brix), day to attain ideal leaf stage for flowering (≤185.70 days), crown weight (≤0.31 kg), peel weight (≤0.16 kg) and acidity (≤1.05). Agronomic information on the selected hybrids is presented in Table 7.

**Table 6.** Average heterosis for the plant and fruit traits used for calculating the value of selection criterion in Mauritius x Kew hybrids.

| Treatment/<br>Hybrid |  | Fruit<br>weight | TSS | Pulp<br>weight | Days to attain<br>physiological maturity | Crown<br>weight | Peel<br>weight | Eye<br>profile | Eye relative<br>surface | Acidity | Value of selection<br>criterion |
| --- | --- | --- | --- | --- | --- | --- | --- | --- | --- | --- | --- |
| T 04 | H 17 | 97.21 | 11.81 | 114.21 | -1.59 | 126.02 | 4.96 | -14.77 | -17.21 | -3.95 | **129.77** |
| T 05 | H 16 | -29.30 | -4.42 | -38.97 | -1.05 | 11.54 | -33.81 | 0.45 | 5.60 | 3.95 | **-59.37** |
| T 06 | H 85 | -36.74 | 2.50 | -44.98 | -2.80 | -20.00 | -31.72 | -18.64 | -6.65 | 0.56 | **0.03** |
| T 07 | H 91 | 4.19 | 14.74 | -8.80 | -1.15 | 28.13 | 7.06 | 44.77 | 3.48 | 2.82 | **-74.98** |
| T 08 | H 48 | 64.65 | 1.65 | 45.96 | -2.49 | 169.92 | 30.31 | 25.45 | 3.70 | -6.21 | **-108.42** |
| T 09 | H 62 | 30.23 | 4.58 | 8.27 | -4.07 | 66.83 | 63.61 | -15.23 | 28.19 | 18.64 | **-114.89** |
| T 10 | H 43 | 39.53 | 21.20 | 45.74 | -0.75 | 56.10 | -22.59 | -18.18 | 4.96 | -8.47 | **95.40** |
| T 11 | H 66 | 34.88 | 7.50 | 48.27 | 0.75 | -25.20 | 0.81 | -12.95 | -24.60 | -2.82 | **154.66** |
| T 12 | H 77 | -36.74 | 5.35 | -66.90 | -1.79 | 80.16 | -44.66 | 4.55 | 5.60 | -0.56 | **-141.59** |
| T 13 | H 92 | -27.44 | 4.89 | -38.33 | 2.07 | -5.04 | -37.51 | 9.09 | 5.60 | 12.99 | **-48.08** |
| T 14 | H 63 | -1.40 | -6.43 | 6.42 | -1.18 | -31.38 | 2.10 | 43.86 | -15.52 | 5.08 | **-4.37** |
| T 15 | H 27 | -18.14 | 0.58 | -47.15 | -4.07 | 126.99 | -29.76 | 16.59 | -5.60 | 0.56 | **-169.42** |
| T 16 | H 78 | 4.19 | 1.89 | -19.40 | -1.32 | 124.72 | -16.84 | 13.64 | 5.60 | 14.12 | **-153.24** |
| T 17 | H 70 | 70.23 | 3.12 | 76.61 | 2.64 | 65.85 | 23.52 | -12.27 | -8.98 | -5.08 | **84.28** |
| T 18 | H 59 | 33.95 | 9.97 | 44.22 | 1.29 | -3.41 | -7.96 | -11.14 | -17.42 | -1.69 | **128.47** |
| T 19 | H 60 | -42.33 | 17.58 | -69.26 | 1.55 | 82.76 | -38.34 | 13.64 | 5.60 | -1.69 | **-157.53** |
| T 20 | H 49 | -24.65 | 13.81 | -38.72 | -0.95 | 19.67 | -29.68 | 25.68 | -5.60 | 8.47 | **-67.15** |
| T 21 | H 54 | 36.74 | 7.12 | 0.73 | -2.33 | 234.31 | -1.79 | 25.68 | -5.60 | 1.69 | **-207.37** |
| T 22 | H 10 | -30.23 | 12.74 | -51.79 | -1.67 | 77.24 | -42.33 | -10.68 | 1.37 | 2.82 | **-96.03** |
| T 23 | H 15 | -33.95 | 5.35 | -55.18 | -1.58 | 60.98 | -32.26 | 13.64 | 5.60 | -0.56 | **-129.60** |
| T 24 | H 30 | -46.98 | 0.65 | -58.94 | 1.65 | -35.61 | -13.45 | -2.95 | 9.82 | 11.86 | **-76.59** |
| T 25 | H 14 | -3.26 | 11.12 | -26.31 | -3.84 | 101.95 | -41.99 | 28.86 | -8.55 | 9.6 | **-104.48** |
| T 26 | H 7 | -6.05 | 7.04 | -22.50 | 0.16 | 40.81 | -14.32 | -23.18 | 0.32 | 16.38 | **-41.68** |
| T 27 | H 35 | 100.00 | 15.97 | 74.18 | -2.04 | 116.19 | 25.05 | -10.68 | -2.64 | -2.82 | **67.09** |
| T 28 | H 19 | -11.63 | 12.89 | -19.26 | -1.73 | 20.96 | -18.78 | -7.50 | -11.09 | 0.56 | **-0.42** |

**Table 7.** Agronomic information on the selected hybrids.

| **Sl. No.** | **Trait** | **H66** | **H17** | **H59** | **H43** | **H70** | **H35** |
| --- | --- | --- | --- | --- | --- | --- | --- |
| 1 | Plant height (cm) | 90.64 | 92.97 | 92.24 | 94.40 | 89.47 | 91.64 |
| 2 | Number of leaves per plant | 42.24 | 40.97 | 41.77 | 38.74 | 41.47 | 41.74 |
| 3 | Length of D leaf (cm) | 58.20 | 54.04 | 61.30 | 56.60 | 58.07 | 55.30 |
| 4 | Breadth of D leaf (cm) | 3.40 | 2.54 | 3.32 | 3.99 | 3.64 | 3.57 |
| 5 | D leaf area (cm<sup>2</sup>) | 143.45 | 99.37 | 147.34 | 163.71 | 153.11 | 142.87 |
| 6 | Leaf Area Index | 4.49 | 3.02 | 4.56 | 4.71 | 4.71 | 4.42 |
| 7 | Days to attain ideal leaf stage for<br>flowering | 181.08 | 182.75 | 178.14 | 184.30 | 190.61 | 181.92 |
| 8 | Days for initiation of flowering<br>(visual) | 47.36 | 46.98 | 47.80 | 49.98 | 45.41 | 45.54 |
| 9 | Days for 50 % flowering | 52.81 | 52.19 | 53.08 | 54.99 | 50.60 | 50.75 |
| 10 | Flowering phase (days) | 21.78 | 20.82 | 21.10 | 20.05 | 20.77 | 20.82 |
| 11 | Length of the fruit (cm) | 17.50 | 16.10 | 14.90 | 13.60 | 15.25 | 10.55 |
| 12 | Girth of the fruit (cm) | 38.50 | 41.10 | 37.80 | 40.80 | 57.40 | 43.20 |
| 13 | Breadth of the fruit (cm) | 11.45 | 11.65 | 11.20 | 12.15 | 9.42 | 9.62 |
| 14 | Crown weight (kg) | 0.42 | 0.35 | 0.26 | 0.24 | 0.26 | 0.33 |
| 15 | Taper ratio of the fruit | 0.82 | 0.91 | 0.84 | 0.89 | 0.93 | 0.89 |
| 16 | Peel weight (kg) | 0.19 | 0.15 | 0.24 | 0.11 | 0.18 | 0.18 |
| 17 | Pulp weight (kg) | 1.08 | 1.59 | 0.80 | 1.08 | 1.31 | 1.29 |
| 18 | Pulp percentage | 79.76 | 89.56 | 70.27 | 85.59 | 83.26 | 70.84 |
| 19 | Days for fruit maturity (days) | 142.82 | 147.07 | 148.24 | 140.33 | 143.02 | 145.77 |
| 20 | Crop duration (days) | 371.25 | 376.79 | 374.17 | 374.61 | 379.04 | 373.22 |
| 21 | Shelf life (days) | 7.38 | 7.50 | 9.00 | 6.84 | 7.71 | 7.34 |
| 22 | Juice (%) | 83.78 | 77.61 | 87.61 | 80.93 | 84.83 | 85.03 |
| 23 | TSS (° Brix) | 13.21 | 14.53 | 13.59 | 15.75 | 13.40 | 15.07 |
| 24 | Acidity (%) | 0.83 | 0.85 | 1.05 | 0.81 | 0.84 | 0.86 |
| 25 | Total sugars (%) | 12.03 | 11.73 | 11.66 | 11.07 | 10.65 | 12.78 |
| 26 | Reducing sugars (%) | 2.61 | 3.65 | 3.11 | 3.53 | 2.98 | 1.92 |
| 27 | Non-reducing sugars (%) | 9.43 | 0.97 | 8.55 | 7.55 | 7.67 | 10.87 |
| 28 | Sugar/ acid ratio | 15.92 | 17.09 | 12.94 | 19.44 | 15.95 | 17.52 |
| 29 | Fibre (%) | 34.03 | 33.28 | 32.80 | 33.53 | 34.26 | 33.15 |
| 30 | Total carotenoids (%) | 267.81 | 275.80 | 240.46 | 278.90 | 325.34 | 238.65 |
| 31 | Ascorbic acid (mg/ 100 g) | 78.64 | 49.40 | 104.11 | 52.31 | 68.55 | 73.34 |
| 32 | Fruit weight with crown (kg) | 1.77 | 2.12 | 1.40 | 1.50 | 1.83 | 2.15 |
| 33 | Profile of eyes | 5.52 | 3.75 | 3.73 | 3.60 | 3.86 | 3.93 |
| 34 | Relative surface of eyes | 4.91 | 3.92 | 6.07 | 4.97 | 4.31 | 4.61 |

Hybridization in pineapple is reported to result in superior hybrid varieties suited for various purposes (Benega et al., 1999). This has resulted hybrids with wide variability in fruit and plant traits, suggesting a high level heterozygosity in the parents used. Heterozygosity has been increased in the present day cultivars, due to many reason but mainly through the self-incompatibility and outcrossing (Sanewski, 2018). Even with the existence of crossing and hybrid variety development, a precise report on the extent of heterobeltiosis, standard heterosis and average heterosis in each trait in pineapple has been missing. In addition to presenting the achievable heterosis targets for each trait in this crop and demonstrating a workable selection criterion, this paper reports the recovery of six promising hybrids, which have to be forwarded for multi-location evaluation and commercial cultivation. The identified hybrids possess significant heterobeltiosis, standard heterosis over the check varieties and average heterosis for nearly all the desirable traits such as fruit size, pulp weight, pulp percentage, TSS, sugar/acid ratio, carotenoid content and ascorbic acid content. Similarly, negative heterosis was observed for most of the undesirable traits.

## Conclusion

Twenty five selected hybrids resulted from the cross of Mauritius and Kew pineapple cultivars were evaluated under RBD in open field conditions. Heterobeltiosis, average heterosis and standard heterosis over two check varieties were worked out in each hybrid, for ten plant growth traits and twenty four fruit traits. A selection criterion, based on the average heterosis of desirable and undesirable traits was formulated and used to identify the promising hybrids. High heterobeltiosis, average heterosis and standard heterosis were observed for most of the desirable traits in the hybrids evaluated and six promising hybrids were recommended from this study.

## Acknowledgments

LD acknowledges Ministry of Tribal Affairs, Government of India, for the National Fellowship for Higher Education of ST Students 2017-2018, for the doctoral research.

## References

Achigan-Dako, E.G., Adjé, C.A., N’Danikou, S., Hotegni, N.V.F., Agbangla, C., and Adam Ahanchédé, A., 2014. Drivers of conservation and utilization of pineapple genetic resources in Benin. SpringerPlus 3: 273. doi: 10.1186/2193-1801-3-273

Adje, C.A.O., Achigan-Dako, E.G., d’Eeckenbrugge, G.C., Yedomonhan, H., and Agbangla, C., 2019. Morphological characterization of pineapple (*Ananas comosus*) genetic resources from Benin. Fruits 74(4): 167–179. doi: 10.17660/th2019/74.4.3

AOAC, 2000. Official Methods of Analysis of AOAC International, Association of Official Analytical Chemists, 17^th^ Ed, Gaithersburg, Md. Association of Official Analytical Chemists Intl.

Balakrishnan, S., Aravindakshan, M., and Nair, N.K., 1978. Efficacy of certain growth regulators in inducing flowering in pineapple (*Ananas comosus* L.). Agric. Res. J. Kerala 16: 125–128.

Bartholomew, D.P., 1977. Inflorescence development of pineapple (*Ananas comosus* [L.] Merr.) induced to flower with ethephon. Bot. Gaz. 138(3): 312–320. doi: 10.1086/336930

Benega, R., Cisneros, A., Hidalgo, M., Martínez, J., Arias, E., Arzola, M., Carvajal, C. and Isidrón, M., 1999. Hybridization in pineapple results and strategies to save time for obtaining and releasing new hybrid varieties for growers. Pineapple News 6: 12–14.

Bhowmik, G. and Bhagabat, A., 1975. Self-incompatibility studies in pineapple (*Ananas comosus* L.). Indian Agric. 19: 259–265.

Brat, P., Hoang, L.N.T., Soler, A., Reynes, M. and Brillouet, J.M., 2004. Physicochemical characterization of a new pineapple hybrid (FLHORAN41 Cv.). J. Agric. Food Chem., 52(20): 6170–6177. doi: 10.1021/jf0492621

Brewbaker, J.L. and Gorrez, D.D., 1967. Genetics of self-incompatibility in the monocot genera, *Ananas* (pineapple) and *Gasteria*. Am. J. Bot. 54 (5, Part 1): 611–616. doi: 10.1002/j.1537-2197.1967.tb10684.x

Cabot, C., 1992. Origin, phylogeny and evolution of pineapple species. Fruits 47(1): 25–32.

Cabral, J.R.S., d’Eeckenbrugge, C.G., and de Matos, A.P., 2000. Introduction of selfing in pineapple breeding. Acta Hortic. 529: 165–168. doi: 10.17660/ActaHortic.2000.529.19

Cabral, J.R.S., Ledo, C.D.S., Cardas, R.C., and Junghans, D.T., 2009. Characters variation in pineapple hybrids obtained by different crosses. Rev. Bras. Frutic. 31(4): 1129–1134.

Cabral, J.R.S., Souza, A.D.S., Matos, A.P.D. and Caldas, R.C., 2003. Effects of self-pollination in pineapple cultivars. Rev. Bras. Frutic. 25(1): 184–185.

Chan, Y.K. and Lee, H.K. 1999. Performance of F_1_ pineapple hybrids selected for early fruiting. J. Trop. Agric. Food Sci. 27: 1–8.

Chan, Y.K., 1991. Evaluation of F_1_ populations from a 4×4 diallel in pineapple and estimation of breeding values of parents. MARDI Res. J. 19: 159–168.

Coppens d’Eeckenbrugge, G., Duval, M.F. and Van Miegroet, F., 1992. Fertility and self-incompatibility in the genus Ananas. In: Proc. First International Pineapple Symposium, Honolulu, Hawaii, USA, 2-6 November 1992, pp. 45–52.

d’Eeckenbrugge, C.G., and Marie, F., 1998. Pineapple breeding at CIRAD. II. Evaluation of ‘Scarlett’, a new hybrid for the fresh fruit market, as compared to ‘Smooth Cayenne’. Acta Hortic. 529: 155–164. doi: 10.17660/ActaHortic.2000.529.18

de Souza, V.A.B., Byrne, D.H., and Taylor, J.F., 2000. Predicted breeding values for nine plant and fruit characteristics of 28 peach genotypes. J. Am. Soc. Hortic. Sci. 125(4): 460–465. doi: 10.21273/JASHS.125.4.460

Dhurve, L., Kumar, K.A., Bhaskar, J., Sobhana, A., Francies, R.M. and Mathew, D., 2021. Wide variability among the ‘Mauritius’ somaclones demonstrates somaclonal variation as a promising improvement strategy in pineapple (*Ananas comosus* L.). Plant Cell Tiss. Org. Cult. 145(3): 701–705. doi: 10.1007/s11240-021-02022-5

Ecostatkerala, 2020. Agricultural Statistics 2017-18. Department of Economics and Statistics, Government of Kerala, India, p.17. http://www.ecostat.kerala.gov.in/index.php/agriculture

Firoozabady, E. and Gutterson, N., 2003. Cost-effective *in vitro* propagation methods for pineapple. Plant Cell Rep. 21: 844–850. doi: 10.1007/s00299-003-0577-x

Fisher, R.A. 1936. The use of multiple measurements in taxonomic problems. Ann. Eugen. 7(2): 179–188. doi: 10.1111/j.1469-1809.1936.tb02137.x

Fournier, P., Dubois, C., and Soler, A., 2007. Considerations on growth characteristics of different pineapple varieties in Ivory Coast, Reunion Island and Caribbean Islands. In: VI International Pineapple Symposium, 18–23 November 2007, Joao-Pessoa, Brésil, Montpellier, CIRAD.

Hadiati, S., Yuliati, S., and Soemargono, A., 2011. Evaluation of qualitative and quantitative characters of pineapple hybrids resulted from crossing between Cayenne and Queen. J. Agric. Bio. Sci. 6(1): 32–38.

Hassan, A., Othman, Z., and Siriphanich, J., 2011. Pineapple (*Ananas comosus* L. Merr.). In: Postharvest Biology and Technology of Tropical and Subtropical Fruits, Mangosteen to White Sapote, Yahia, E.M. (Ed.) ISBN 978-0-85709-090-4, Woodhead Publishing, pp 194–217, 218e. doi:10.1533/9780857092618.194

IBPGR, 1991. Descriptors for Pineapple, International Board for Plant Genetic Resources, Rome, Italy, ISBN 978-92-9043-199-2, 41p.

KAU, 2016. Package of Practices Recommendations – Crops, 15 Ed., Kerala Agricultural University, Thrissur, India, p. 223–226. Available at: https://kau.in/document/11029

Kishore, K., Rupa, T.R. and Samant, D., 2021. Influence of shade intensity on growth, biomass allocation, yield and quality of pineapple in mango-based intercropping system. Sci. Hortic. 278: 109868. doi: 10.1016/j.scienta.2020.109868

Kuriakose, K.P. 2004. Heterosis breeding and in vitro mutagenesis in pineapple (Ananas comosus [L.] Merr.). Ph. D. thesis in Horticulture, Kerala Agricultural University, Thrissur, India, 212 p.

Laufer, B. 1929. The American plant migration. Sci. Monthly 28: 239–259.

Marie, F., d’Eeckenbrugge, C.G., and Bernasconi, B., 1998. Pineapple breeding at CIRAD I. Evaluation and selection of ‘Smooth Cayenne’ x ‘Manzana’ hybrids. Acta Hortic. 529: 147–154. doi: 10.17660/ActaHortic.2000.529.17

Moreira, S.O., Kuhlcamp, K.T., Barros, F.L.D.S., Zucoloto, M., and Godinho, T.D.O., 2019. Selection index based on phenotypic and genotypic values predicted by REML/BLUP in papaya. Rev. Bras. Frutic. 41(1): e-079. doi: 10.1590/0100-29452019079

Paull, R.E., Bartholomew, D.P., and Chen, C.C., 2017. Pineapple breeding and production practices. In: Handbook of Pineapple Technology: Production, Postharvest Science, Processing and Nutrition, Eds. Lobo, M.G. and Paull, R.E., ISBN: 9781118967386, John Wiley & Sons, Ltd., West Sussex, UK, pp.16–38. doi: 10.1002/9781118967355.ch2

Rasmusson, D.C., 1987. An evaluation of ideotype breeding. Crop Sci. 27: 1140–1146. doi: 10.2135/cropsci1987.0011183X002700060011x

Sanewski, G.M., 1998. The Australian pineapple fresh market breeding program. In: Abstracts of Third International Symposium, 17–20 November 1998, Department of Agriculture, Pattaya, Thailand, p. 51.

Sanewski, G.M., 2009. The effect of different levels of inbreeding on self-incompatibility and inbreeding depression in pineapple. Acta Hortic. 822: 63–70. doi: 10.17660/ActaHortic.2009.822.6

Sanewski, G.M., 2018. The history of pineapple improvement. In: Genetics and Genomics of Pineapple, Plant Genetics and Genomics: Crops and Models 22, Ed. Ming, R., p. 87-96. doi: 10.1007/978-3-030-00614-3_7

Sen, S.K., 2001. Pineapple. In: Fruits of India: Tropical and Subtropical, Ed. Bose, T.K., Naya Prakash, Culcutta.

Sheoran, O.P., Tonk, D.S., Kaushik, L.S., Hasija, R.C., and Pannu, R.S., 1998. Statistical Software Package for Agricultural Research Workers. In: Recent Advances in Information Theory, Statistics and Computer Applications, Eds., Hooda, D.S. and Hasija, R.C., CCS Haryana Agricultural University, Hisar, India, p. 139–143. Available at http://14.139.232.166/opstat/

Souza, F.V.D., Cabral, J.R. dos S. de Souza, E.H. Silva, M.de J., Santos, O.S.N., and Ferreira, F.R. 2009. Evaluation of F_1_ hybrids between *Ananas comosus* var. *ananassoides* and *Ananas comosus* var. *erectifolius*. Acta Hortic. 822: 79–84. doi: 10.17660/ActaHortic.2009.822.8

Van de Poel, B., Ceusters, J., and De Proft, M.P. 2009. Determination of pineapple (Ananas comosus, MD-2 hybrid cultivar) plant maturity, the efficiency of flowering induction agents and the use of activated carbon. Sci. Hortic. 120(1): 58–63. doi: 10.1016/j.scienta.2008.09.014

Van Overbeek, J. and Cruzado, H.J., 1948. Note on flower formation in the pineapple induced by low night temperatures. Plant Physiol. 23(3): 282–285. doi: 10.1104/pp.23.3.282

Viana, E.D.S., Reis, R.C., Jesus, J.L.D., Junghans, D.T., and Souza, F.V.D., 2013. Physico-chemical characterization of new hybrids pineapple resistant to fusariosis. Ciênc. Rural 43(7): 1155–1161. doi:10.1590/S0103-84782013005000075

Viana, E.D.S., Sasaki, F.F.C., Reis, R.C., Junghans, D.T., Guedes, I.S.A., and Souza, E.G., 2020. Quality of fusariosis-resistant pineapple FRF 632, harvested at different maturity stages. Rev. Caatinga 33(2): 541–549. doi:10.1590/1983-21252020v33n226rc

Wang, R.H., Hsu, Y.M., Bartholomew, D.P., Maruthasalam, S. and Lin, C.H., 2007. Delaying natural flowering in pineapple through foliar application of aviglycine, an inhibitor of ethylene biosynthesis. HortScience 42(5): 1188–1191. doi: 10.21273/HORTSCI.42.5.1188

Watson, D.J., 1952. The physiological basis of variation in yield. In: Advances in Agronomy, Ed. Norman, A.G., Vol. 4, ISBN: 9780080563176, Academic Press, Cambridge, Massachusetts, pp. 101–145.

